# Structure, function and dynamics of mCoral, a pH responsive engineered variant of the mCherry fluorescent protein with improved hydrogen peroxide tolerance

**DOI:** 10.1101/2025.11.26.690718

**Authors:** Athena Zitti, Ozan Aksakal, Danoo Vitsupakorn, Pierre J. Rizkallah, Halina Mikolajek, James A. Platts, Georgina E. Menzies, D. Dafydd Jones

## Abstract

The red fluorescent protein mCherry is one of the most utilised fluorescent proteins in biology. Here, we have changed the chromophore chemistry by converting the thioether group of M66 to a thiol group through mutation to cysteine. The new variant termed mCoral due to its orange fluorescence hue has similar brightness to mCherry but has improved resistance to hydrogen peroxide. The variant is also responsive to pH with a low and high pKa forms that have distinct spectral properties, which DFT analysis suggests is due to protonation state changes in the newly introduced thiol group as well as the phenol group. The structure of mCoral reveals that the M66C mutation creates a space within the β-barrel structure that is filled by a water molecule, which makes new polar interactions including with backbone carbonyl group of F65. Molecular dynamic simulations suggests that this additional water molecule, together with local solvation around the chromophore, could play a role in promoting planarity of the full conjugated system comprising the chromophore; the mCoral chromophore makes slightly more H-bonds with water than mCherry. The main water exit point for mCherry is also narrower in mCoral potentially explaining the increased resistance to hydrogen peroxide. Overall, a small structural change to mCherry has resulted in a new fluorescent protein with potentially useful characteristics and an insight into the role of dynamics and water in defining structure-function relationship in red fluorescent proteins.

## 1. Introduction

Fluorescent proteins (FPs) have revolutionised molecular and cellular biology by allowing genetic tagging of defined targets enabling non-invasive imaging of biological processes ^1–3^. Among the many available FPs, mCherry ^4^ is one of the most widely used. This red FP originally derived from the original *Discosoma* sp. DsRed ^5^ FP is popular due to its relatively simple monomeric structure, brightness and photostability ^4,6^. However, as with many FPs, mCherry’s fluorescence is influenced by environmental factors, including pH and reactive oxygen species (ROS) such as hydrogen peroxide.

The pH sensitivity of FPs is a critical factor to consider, especially when imaging cellular compartments that have different pH environments to the cytosol ^7^. While the cytosol is normally around pH 7, the Golgi apparatus is lower at pH 6.0-6.7, endosomes and secretory vesicles are lower again at pH 5.0-6.5 while lysosomes are around 4.5-5.0; the mitochondrial matrix samples a slightly higher pH regime (7.5-8.0). While being able to monitor processes in different compartments is important for understanding various cellular processes such as secretion, autophagy, endocytosis and cellular stress, the right FP needs to be selected for the job. While mCherry is thought to be relatively stable to mildly acidic conditions with a reported pKa of 4.5 ^4^, it can be influenced by other factors such fusion partner and local protein concentrations. Furthermore, pH responsiveness of mCherry is not completely understood but is thought to solely involve protonation of the phenolic group of the chromophore (Figure 1a). Given the importance of pH to biology ^8,9^, there has been a concerted effort to develop pH responsive FPs that report on changes in cellular pH; most such as SypHer ^10,11^, mKeima ^12^ and pHlorin ^13–15^ have fluorescence in the green-yellow range. These FPs have some drawbacks such as limited pH range (e.g. either high range or low range), have modest dynamic ranges and can be sensitive to ROS such as hydrogen peroxide.

**Figure 1.**
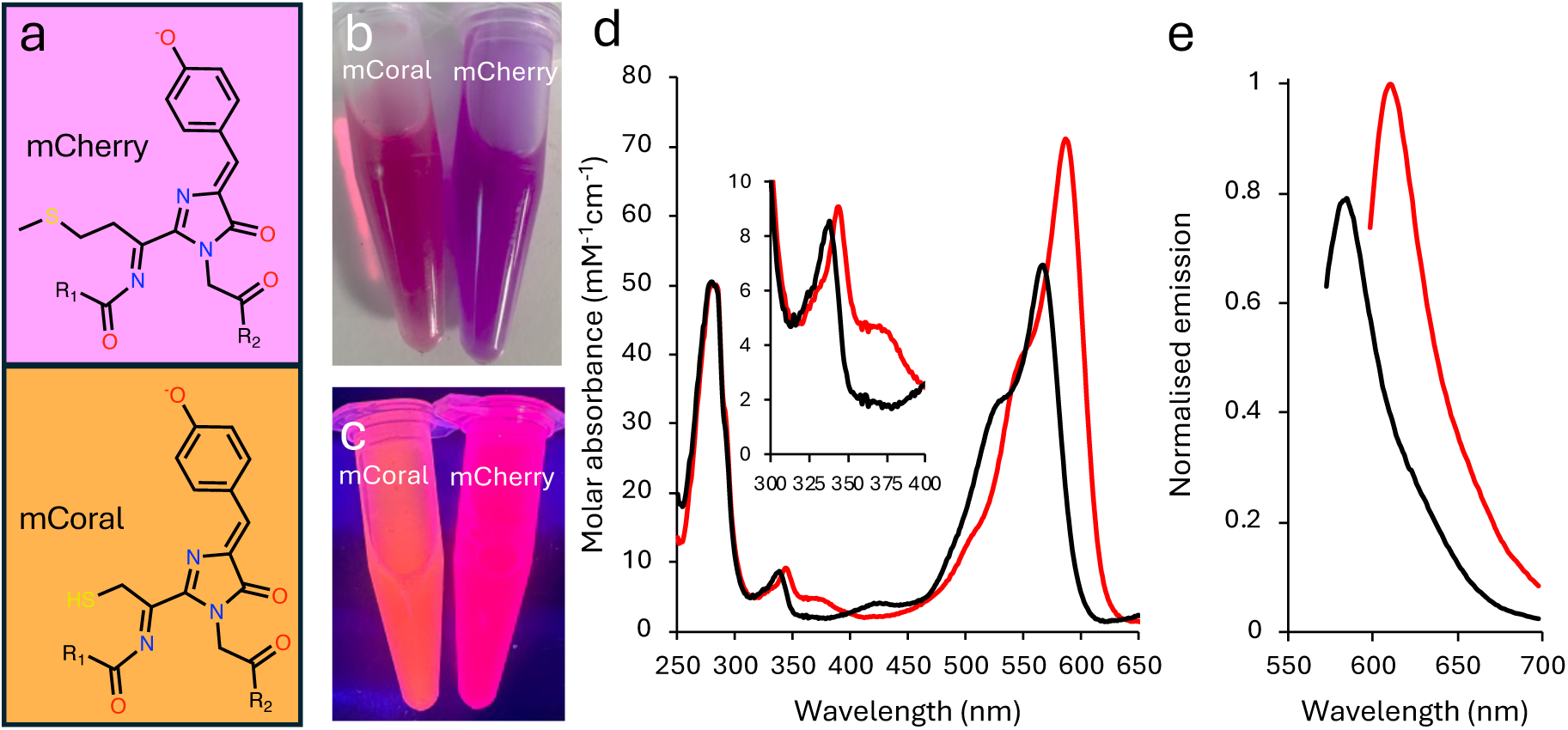
The effect of introducing cysteine into the mCherry chromophore. (a) The chromophore structures of mCherry (66-MYG-68 tripeptide) and the M66C variant (termed mCoral). The protein colour in (b) ambient light and (c) on exposure to UV light. (d) the absorbance spectra of mCherry (red) and mCoral (black). (e) Emission spectra of mCherry (red) and mCoral (black) on excitation at their λ_max_ wavelengths in (d). Fluorescence emission (5 μM of each protein) normalised to mCherry based their respective λ_EM_ values.

As well as pH flux, ROS like hydrogen peroxide, play an important role in biological systems ^16,17^ and can impact on FP function^18^. Resistance to H_2_O_2_ is particularly important as it is generated as part of the chromophore maturation process ^19–21^ and the chromophores themselves can promote the formation of H_2_O_2_ ^22–24^. One of the design concepts for improved resistance to H_2_O_2_ is the removal of sulphur-containing amino acids such as methionine and cysteine, as sulphur oxidation is a common mechanism of action ^25^. H_2_O_2_ is also an important biological signalling molecule so FPs have been generated detected changes in its level. These sensors such as the HyPer series ^26,27^ are normally based on a domain insert system, with a H_2_O_2_ responsive protein such as OxyR inserted within a FP.

Here, we have taken the commonly used red fluorescent protein, mCherry ^4,28^ and converted it into a pH responsive variant with improved resistance to H_2_O_2_. This new variant, termed mCoral, has a single mutation, M66C, within the chromophore forming amino acid XYG sequence. Structure determination reveals that mCoral is similar to mCherry but its chromophore has a different hydration pattern. Molecular dynamic simulations suggest that the additional water molecule occupying the space left on side chain shortening at residue 66 helps retain the extended chromophore planar state of the chromophore. Also, the main water exit point is narrower in mCoral providing an explanation for improved resistance to H_2_O_2_.

## 2. Results and discussion

### 2.1 Generation of mCoral

The chromophore of typical β-barrel fluorescent proteins is comprised of the tripeptide sequence XYG, where the first amino acid is variable ^3,29,30^. The CYG sequence is relatively uncommon as a chromophore forming sequence, with less than 5% of FPs in FPBase ^31^ containing this tripeptide sequence; many of these derive from *Echinophyllia Sp* 22G dronpa-based photoswitchable variants ^32^. The majority of CYG chromophores also excite below 550 nm. Incorporation of cysteine within the chromophore does however open up the potential for new chemical properties not available to the other 19 common natural amino acids that can dynamically tune the spectral characteristics of a FP. These include redox activity ^33,34^, a physiologically relevant pKa (∼8) ^35^, metal ion binding ^36^ and covalent modification ^37^. We thus sought to introduce cysteine into the commonly used red FP, mCherry ^4,28^ via the M66C mutation to form the chromophore outlined in Figure 1a.

Introduction of the M66C mutation results in a functional, fluorescent protein with a change in inherent observable colour (Figure 1b-c). In ambient light, mCherry turns from a dark purple to burgundy on incorporating the M66C mutaiton (Figure 1b); when irradiated with UV light, the M66C mutation results in mCherry turning from a pink to an orange coral colour (Figure 1c). For this reason, we now term the mCherry M66C variant mCoral. Compared to mCherry, the absorbance and emission spectra are blue shifted (Figure 1d-e and Table 1). The mCoral absorbance spectra has the characteristic double hump absorbance spectrum of mCherry (Figure 1d) but with λ_max_ shifted by 21 nm and the molar absorbance 0.73 fold that of mCherry (Table 1 and Figure 1d). The brightness of mCoral is similar to mCherry due to an improved quantum yield (Table 1).

**Table 1.**
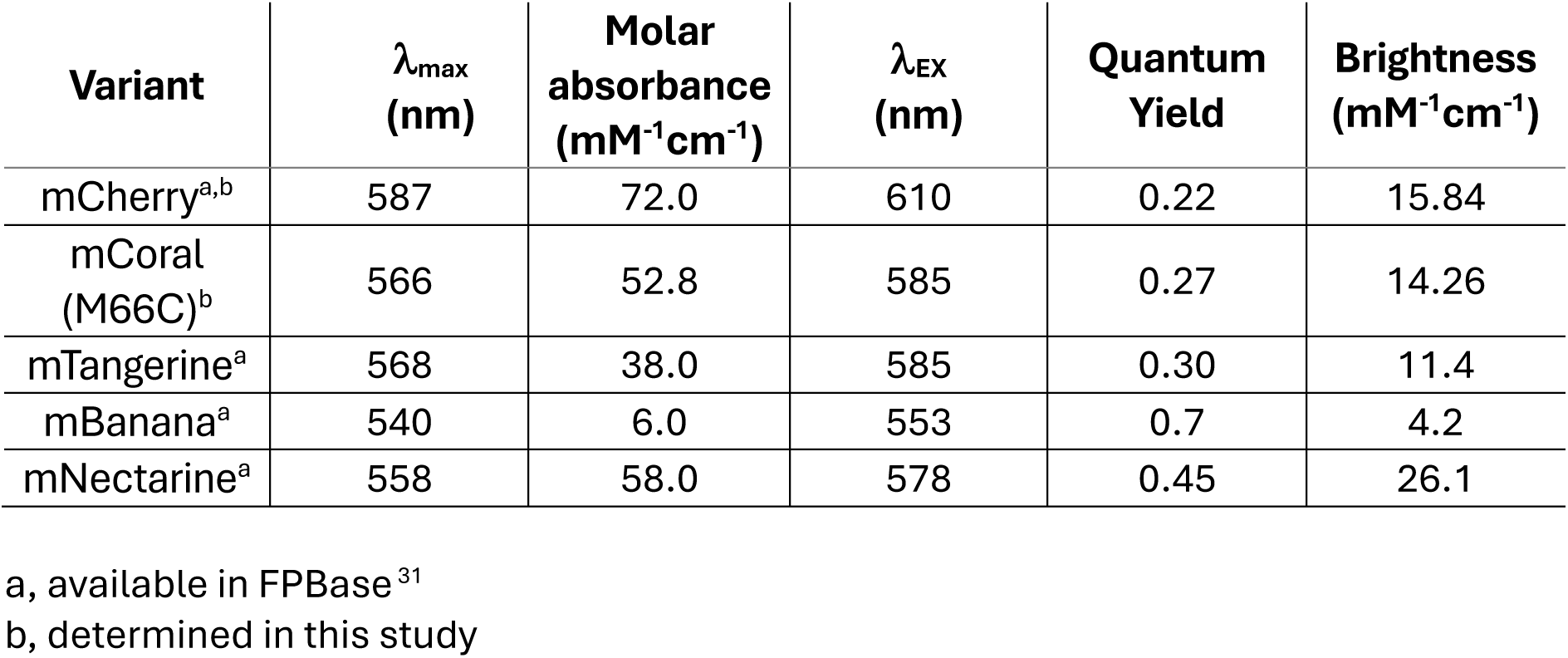
Spectral properties of selective RFPs.

| Variant | $\lambda_{\max}$<br>(nm) | Molar<br>absorbance<br>(mM<sup>-1</sup>cm<sup>-1</sup>) | $\lambda_{\text{EX}}$<br>(nm) | Quantum<br>Yield | Brightness<br>(mM<sup>-1</sup>cm<sup>-1</sup>) |
| --- | --- | --- | --- | --- | --- |
| mCherry<sup>a,b</sup> | 587 | 72.0 | 610 | 0.22 | 15.84 |
| mCoral<br>(M66C)<sup>b</sup> | 566 | 52.8 | 585 | 0.27 | 14.26 |
| mTangerine<sup>a</sup> | 568 | 38.0 | 585 | 0.30 | 11.4 |
| mBanana<sup>a</sup> | 540 | 6.0 | 553 | 0.7 | 4.2 |
| mNectarine<sup>a</sup> | 558 | 58.0 | 578 | 0.45 | 26.1 |

a, available in FPBase<sup>31</sup>
b, determined in this study

Three other DsRed derived FPs contain the CYG chromophore: mTangerine^4^, mBanana ^4^ and mNectarine ^35^. The mCherry derived mCoral compares favourably to mTangerine and mBanana with higher brightness primary due to improved molar absorbance (Table 1). In contrast the directly evolved mNectarine does still appear to be a better protein in terms of brightness but is blue shifted compared to mCoral ^35^. Furthermore, the reported absorbance spectra at pH 7 for mNectarine is very complex with at least 3 major peaks observed, which is likely due to multiple ionisable chromophores forms being present. In comparison, mCoral has more defined spectral profile at pH 7 (Figure 1d-e).

### 2.2 The eFect of pH on mCoral

We next looked at the effect of pH on spectral properties of mCoral. The spectral properties suggest three distinct species for mCoral. At neutral pH, the major absorbance (and excitation) peak is at 566 nm (Figure 2a-b). At higher pH (>9) the major absorbance (and excitation) peak blue shits to 543 nm with the major emission peak also blue shifted to 562 nm (Figure 2a-b); the absorbance spectrum at pH 8 shows a transition between the two forms. At lower pH (<5) a blue shifted peak absorbance peak dominates at 420 nm (Figure 2a) that does not appear to have any significant fluorescence associated with it (Figure S1). Using the absorbance ratios of the major peaks at each pH, we calculated two pKa for mCoral to be 5.7 and 8.8 (Figure 2c-d). When repeated with mCherry, there is no shift in peak wavelengths only a drop in the peak intensity (Figure S2). A single pH transition is observed for mCherry with a pKa of 4.4 (Figure S2), which is similar to that reported previously (pKa 4.5) ^4^.

**Figure 2.**
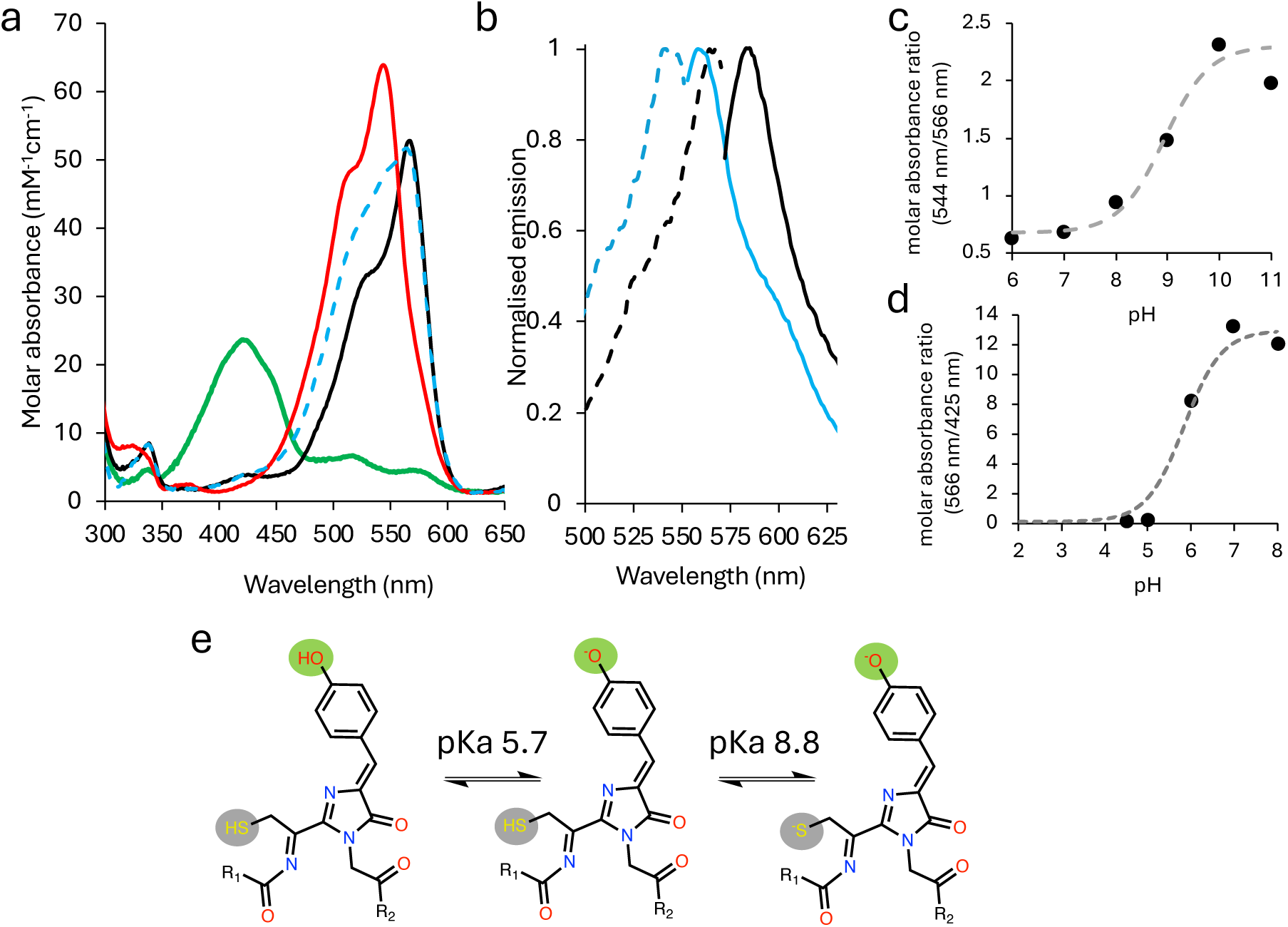
The pH-dependent spectral profile of mCoral. (a) the absorbance spectra of mCoral at pH 4.5 (green), pH 7 (black), pH 8 (dashed blue line) and pH 10 (red). (b) The excitation (dashed lines; emission at 580 nm) and emission spectra (solid lines; on excitation at the variant’s λ_max_) of mCoral at pH 10 (blue) and pH 7 (black). Emission is normalised to the pH 7 values. (c) Ratio plot of absorbance at 544 nm and 566 nm at pH 6 to 11 to determine high range pKa. (d) Ratio plot of absorbance at 566 nm and 425 nm at pH 4.5 to 8 to determine low range pKa. (e) Proposed model for different ionised states of mCoral chromophore giving rise to the different spectral forms. The pKa values were calculated based on the plots in (c) and (d).

To try and understand the basis for the pH effect on mCoral we performed density function theory (DFT) calculations on each of the ionised states of the chromophore using the known chromophore structure (vide infra for mCoral). The chromophore is isolated from the crystal structure and capped as amides in the position of flanking residues. Each possible protonation state is constructed, fully geometry optimised and absorption spectrum predicted. When we compare the DFT predicted absorbance values for each of the charged forms of the mCoral chromophore, the low pH form is likely to be protonated C_66_-SH/Y_67_-Ph-OH form (predicted λ_max_ 422nm; Table 2). The high pH form is likely to be the deprotonated thiolate-phenolate (C_66_-S^-^/ Y_67_-Ph-O^-^, predicted λ_max_ 541; Table 2). At neutral pH, two forms are possible: the thiolate-phenolic (C_66_-S^-^/ Y_67_-Ph-OH) or the thiol-phenolate (C_66_-SH/ Y_67_-Ph-O^-^). Given that the pKa for the phenolic-phenolate transition normally lies in the pH 4-6 range coupled with the DFT prediction that major absorbance for C_66_-SH/Ph-O^-^ peaks at 557 nm, we suggest that the neutral pH form is populated largely by the thiol/phenolate chromophore form. The overall predicted pH conversion scheme is shown in Figure 2e. For mCherry, the Ph-OH is predicted to be absorb at 438 nm with relatively weak oscillator strength, which would appear to be the origin of the new low intensity peak observed in the absorbance spectrum at pH 4.5 (Figure S2).

**Table 2.**
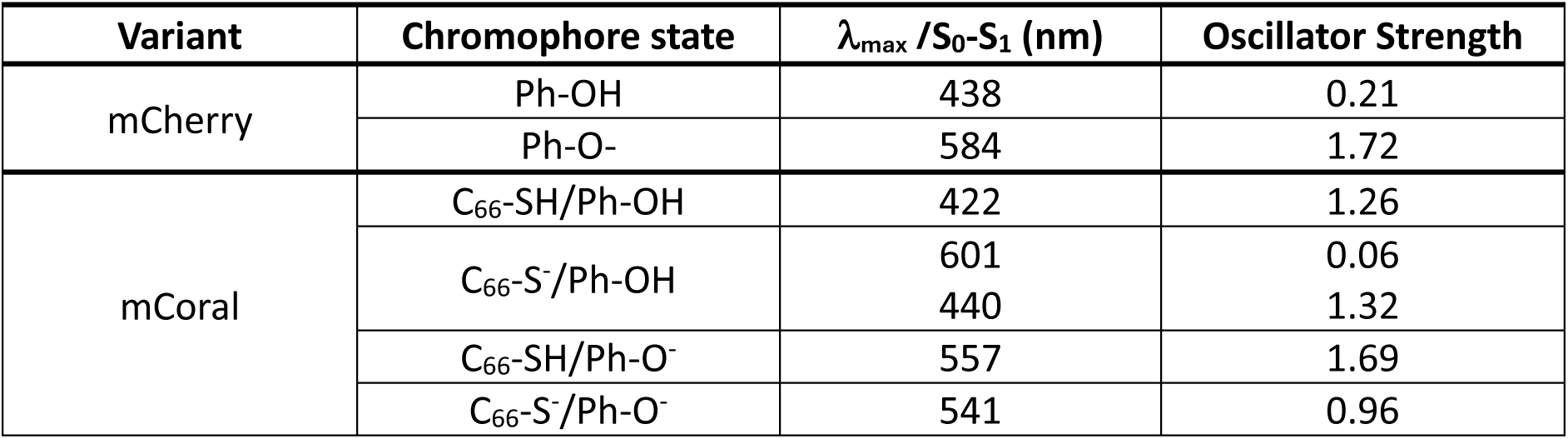
Predicted maximal absorbance wavelengths for each chromophore ionised state.

Insight into the origin of blue/red shifts resulting from pH changes can be drawn from analysis of individual orbitals involved. All strong excitations are dominated by highest occupied molecular orbital (HOMO) to lowest unoccupied molecular orbital (LUMO) excitation, so analysis focuses on these. In the fully protonated form, the HOMO is localised mainly on the phenol and heterocycle, while LUMO lies closer to the cysteine, with energy difference between frontier orbitals ΔE = 3.2 eV. Deprotonating at phenol does not strongly change the localisation of orbitals and hence raises the energy of the HOMO more than that of the LUMO to give ΔE = 2.7 eV, consistent with the red-shift seen at neutral pH. Further deprotonation at SH further raises energy of both HOMO and LUMO, but now affects the LUMO more due to its proximity to cysteine, resulting in ΔE = 2.84 eV and so predicts blue shift compared to SH/O- species.

### 2.3 Influence of H_2_O_2_ on mCoral spectral properties

Sulphur-containing amino acids such as methionine and cysteine can be chemically oxidised by biologically relevant reactive oxygen species (ROS) such as hydrogen peroxide ^38,39^. Thus, the presence of a methionine (mCherry) or cysteine (mCoral) in the chromophore can have its disadvantages and advantages. The disadvantages include loss of fluorescence and thus cellular detection on chemical modification, but this can also be advantageous as such modifications could lead to changes in spectral properties leading to the ability to sense these important ROS species ^33,34,37^.

Both mCherry and mCoral are sensitive to H_2_O_2_ but to different extents. With regards to their spectral properties, mCoral is more stable to H_2_O_2_ compared to mCherry (Figure 3a-c). After 1hr in 0.1% (v/v) H_2_O_2_, mCoral losses less than ∼10% of its fluorescence emission while mCherry’s drops by ∼35% (Figure 3a). The fluorescence emission of mCherry is effectively lost at 1 hr in 0.5% (v/v) H_2_O_2_ while mCoral still retains ∼35% of its signal after 1 hr in 1% (v/v) H_2_O_2_ (Figure 3b). The absorbance spectra confirm there are significant changes to the chromophore on exposure to H_2_O_2_ (Figure 3c). The major absorbance peaks are lost for mCherry suggesting loss of chromophore integrity while mCoral still retains its absorbance peaks, albeit at a lower level (Figure 3c).

**Figure 3.**
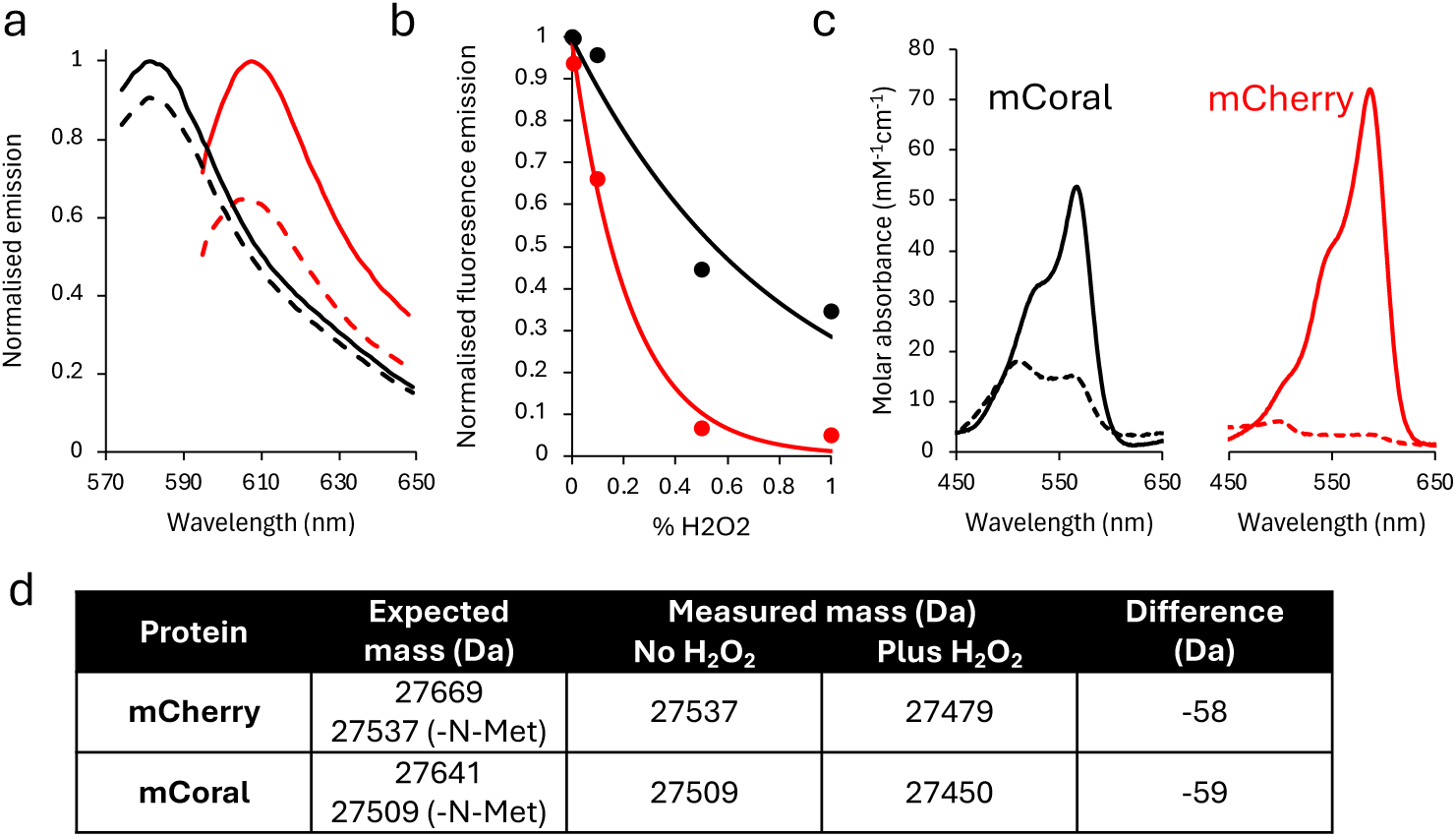
The effect of H_2_O_2_ on the spectral properties of mCoral (black) and mCherry (red). (a) Change in fluorescence emission before (solid lines) and after 1 hr incubation with 0.1% (v/v) H_2_O_2_ (dashed lines). (b) Fluorescence emission on incubation for 1 hr with different concentrations of H_2_O_2_. (c) Absorbance spectra before (solid lines) and after 1 hr 0.5% (v/v) H_2_O_2_ (dashed lines). (d) summary of expected versus observed protein masses before and after the addition of 0.1% (v/v) H_2_O_2_. Mass spectra can be found in Supplementary Figure S3

To understand the chemical modification process likely to be driving the spectral changes, we determine the mass before and after H_2_O_2_ addition (Figure 3d and Figure S3). For both proteins, when taking into account N-terminal methionine removal and chromophore maturation, the expected molecular mass is observed prior to the addition of H_2_O_2_. On addition of H_2_O_2_, both lose 58-59 Da in mass. We currently do not know what process is occurring for both proteins to lose a similar amount of mass, especially as the most common oxidation event involving sulphur-containing amino acids is oxygenation of the sulphur atom ^38,39^, which would increase the mass in units of 15-16 Da. Given that the loss in mass is the same for both proteins, we suggest that a common chemical event is occurring but is either occurring quicker in mCherry or has less of an impact in mCoral, based on the effect of H_2_O_2_ in spectral properties (Figure 3a-b).

### 2.4 Structure and dynamics of mCoral

To understand how replacing methionine for cysteine in the mCherry chromophore affects spectral properties, we determined the structure mCoral (see Table S1 for statistics). Overall, the M66C mutation does not change the general protein structure compared to mCherry (Figure 4a), with the Cα root mean square deviation (RMSD) being 0.152 Å over the whole protein. The chromophore and local protein environment surrounding the mutation site is also largely similar (Figure 4b). Water molecules (W1 to W4 in Figure 4c) retain similar positions to that observed in mCherry. The loss of -SCH_3_ group on conversion of methionine to cysteine will result in space becoming available for an additional moiety; based on obversed electron density we have modelled a water (W_M66C_ in Figure 4d) to take the place of the original M66 side chain component. This additional water makes a new set of polar contacts with backbone carbonyl of F65, the new thiol group in the chromophore and the sidechain carboxamide group of Q42. This new water molecule may also help stabilise the negative charge on the thiolate at high pH as there are no basic residues close by. When we assess potential tunnels through to the new thiol group, access points calculated by CAVER ^41^ are found to be relatively distant from the chromophore, towards the two ends of the β-barrel structure (Figure 4e). One tunnel (coloured red in Figure 4e) exits between stand S10 and the region bisecting strand 7 comprising residues 140 to 143 which is a bulge type loop in the crystal structure.

**Figure 4.**
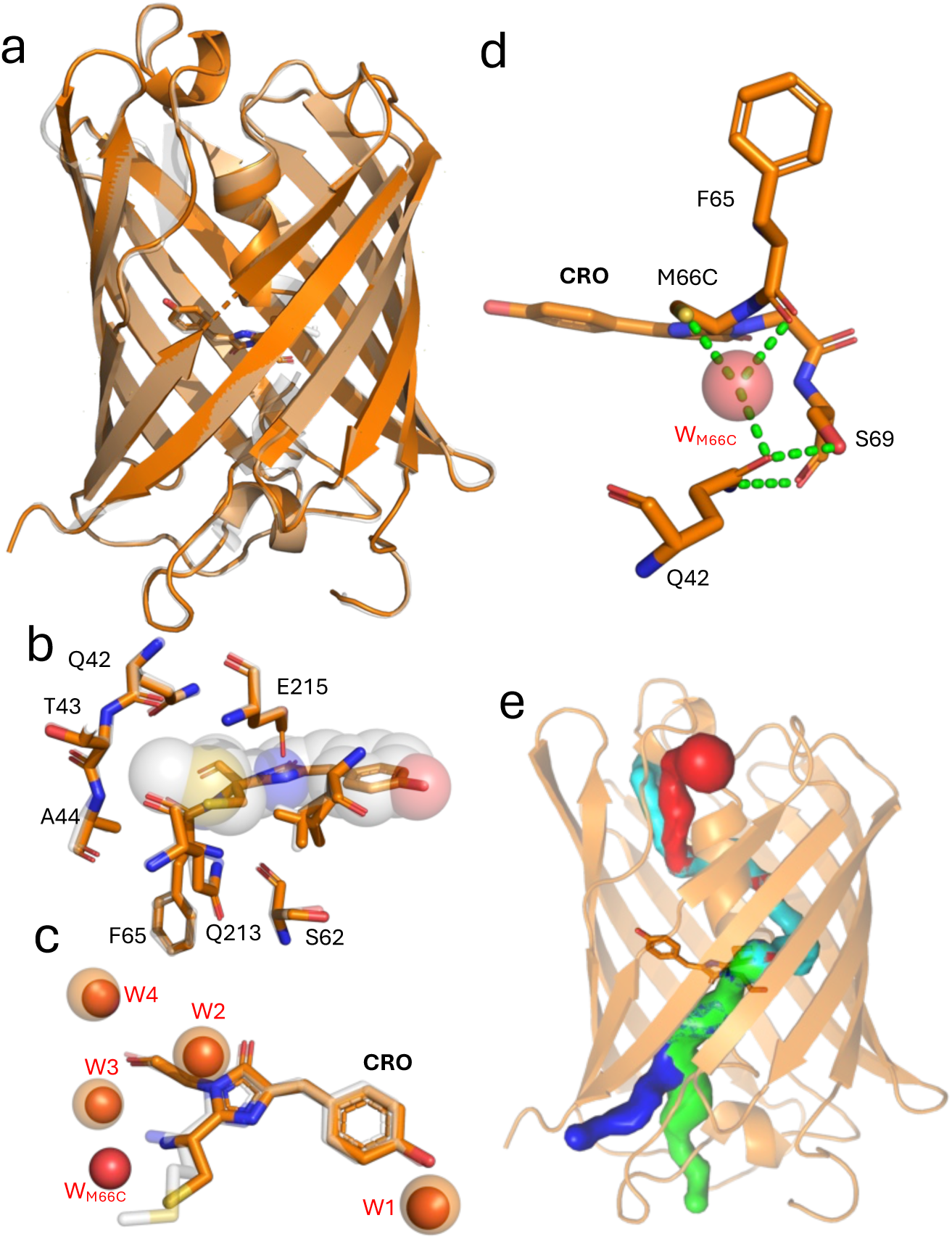
Structure of mCoral. (a) Structural alignment of mCoral (orange) and mCherry (grey; PDB 2h5q ^28^). The chromophore is shown as sticks. (b) Structural alignment of mCoral local chromophore environment (orange) with that of mCherry (grey). The mCherry chromophore is shown as spheres. (c) Local water molecules surrounding chromophore in mCherry (grey with waters shown as transparent spheres) and mCoral (orange with waters shown as solid red spheres). (d) Polar network (shown as dashed green lines) involving water W_M66C_ (red sphere) calculated using the PyMOL ^40^ polar contacts tool. (e) Calculated water tunnels into the void resulting from the M66C mutation. Tunnels (coloured blue, green, cyan and red) were calculated using CAVER3.0.^41^

We then turned to molecular dynamics (MD) to provide further insight into mCoral’s structure-function relationship. As has been applied previously ^42^, our MD simulations use a chromophore that includes F65 to incorporate the additional double bond formed on maturation (see Figure S4). The overall average Cα RMSD over 3×1000 ns production MD runs for mCoral is relatively stable (Figure S5) with an average RMSD of 0.13 ns ± 0.01, slightly lower than mCherry (0.16 ns ± 0.01). The per reside Cα root mean square fluctuation (RMSF) remains similar except for the CRO and its adjacent residues (Figure 5a). The mCherry CRO has a higher flux (0.11 nm ± 0.04) over the simulations compared to mCoral (0.051 nm ± 0.01).

**Figure 5.**
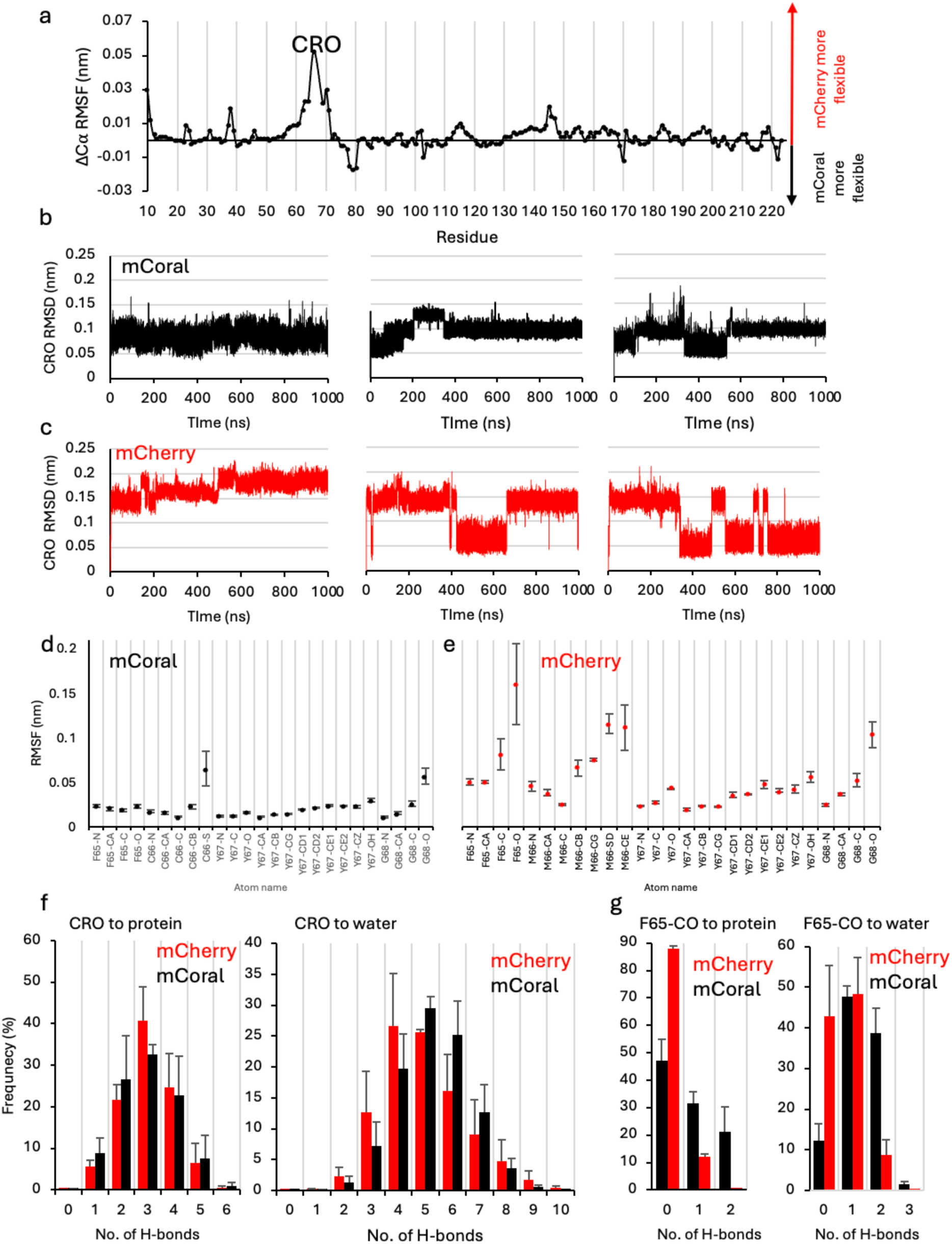
Molecular dynamics analysis of mCherry (red throughout) and mCoral (black throughout). (a) Difference RMSF calculated by subtracting the mCoral residue Cα from the same residue of mCherry. Positive values represent a higher RMSF for mCherry and negative values a higher RMSF for mCoral. The individual RMSF for each variant is shown in Figure S6. The CRO RMSD of the (b) mCoral and (c) mCherry. The per heavy atom CRO RMSF for (d) mCoral and (e) mCherry. The atom nomenclature is shown in Suppering Figure S7. (f) The number of H-bonds frequency between the CRO and either the rest of the protein or water. (g) Number of H-bond frequency between the carbonyl group of F65 and either the rest of the protein or water.

The probe the CRO dynamics further, the RMSD of the chromophore (all atoms) was measured and found to differ between mCherry and mCoral (Figure 5b-c). RMSD analysis suggests that the mCherry CRO is switching between at least two distinct conformations. In comparison, mCoral’s CRO appears more stable with some alternative states observed but with RMSD not switching to the extent observed for mCherry. To identify the CRO atoms contributing the step changes in RMSD, atom-by-atom RMSF was performed on the CRO (Figure 4d-e). For mCoral, the fluctuation of each atom was low (average of 0.022 nm ± 0.012), with the thiol sulphur of the newly introduced C66 (C66-S in Figure 5d) being most dynamic atom. In contrast, mCherry displayed higher per-atom flux (0.054 nm ± 0.034). More specifically, the carbonyl oxygen from F65 (F65-O in Figure 5d-e) is highly dynamic in mCherry but is relatively stable in mCoral. As mentioned above, the F65 carbonyl oxygen forms interactions with the newly observed W_M66C_ water molecule in mCoral (Figure 4d). This could suggest an important role for the W_M66C_ water molecule in stabilising the chromophore conformation. As with mCoral, the side chain of the residue 66 component of the mCherry chromophore is also more dynamic than most of the chromophore heavy atoms (Figure 5e). This suggests despite being packed within the core of the protein, rotation of the -SCH_3_ group is possible in mCherry.

Given the potential importance of local polar interactions between the CRO and its protein and solvent environment (see Figure 4c-d), we measured the number and persistence of H-bonding involving the CRO in each variant. Throughout the simulations, the CRO makes extensive H-bonds with both the rest of the protein and water (Figure 5f), with an average of over 1.2×10^6^ observed over 1 μs (100001 frames) MD simulation for each protein. The mCherry CRO makes slightly more H-bonds with the rest of protein (average of 4.6×10^5^; 4.6 per frame) compared to mCoral (average of 4.4×10^5^; 4.4 per frame). Both have a median value of 3 H-bonds to the rest of the protein but the frequency of 3 or more H-bonds is higher for mCherry (Figure 5f). In contrast, mCoral makes slightly more H-bonds with water molecules (average of 7.9×10^5^ versus 7.5×10^5^; 7.9 per frame versus 7.5 per frame). The number of H-bonds with water spans a greater range with the highest frequency interactions in mCoral at a higher H-bond count (5-6 H-bonds per frame) compared to mCherry (4-5 H-bonds per frame). When the carbonyl oxygen of F65 is assessed specifically, H-bonds are formed more frequently with both protein and water in mCoral (Figure 5g). On average the mCoral carbonyl oxygen of F65 forms 6-fold and 2-fold more H-bonds with either the rest of the protein or water, respectively, than in mCherry. Thus, introduction of the M66C mutation coupled with the incorporation of a new water molecule leads changes in the local chromophore interaction network.

Isolation of individual trajectories representative of the different observed RMSD states shows that the F65 carbonyl oxygen flips position during the mCherry simulation (Figure 6a). In the high RMSD form, the carbonyl oxygen flips ∼180° compared the lower RMSD form representative of the crystal structure conformation resulting in the last double bond being out of plane with the rest of the conjugated network. Linked to the flip is the configuration of M66 side chain which changes relative positions of the Cβ, Cχ and S atoms (Figure 6a). With respect to mCoral, there is rotation of the F65 carbonyl oxygen (up to 50°) but within the plane of the conjugated bond system (Figure 6b). The thiol sulphur also changes it positioning. DFT calculations performed on the alternate chromophore configuration identified for mCherry predicts that the protonated form will absorb at 369 nm (strength 1.18), and the deprotonated form at 422 nm (strength 0.90), representing significant blue-shift compared to the dominant conformation. There is a small absorbance peak at ∼380 nm only present in the mCherry spectrum (Figure 1d, inset). However, given the frequency observed for the alternative mCherry CRO state in the MD simulations, it may not be sampled as frequently or be as persistent in the protein in solution resulting in minimal changes to steady state spectral properties. The MD simulations suggest that the M66C mutation stabilises the conformation of the chromophore, with the local water interactions likely to be playing a key role. This may be the reason for the improve quantum yield of mCoral compared to mCherry.

**Figure 6.**
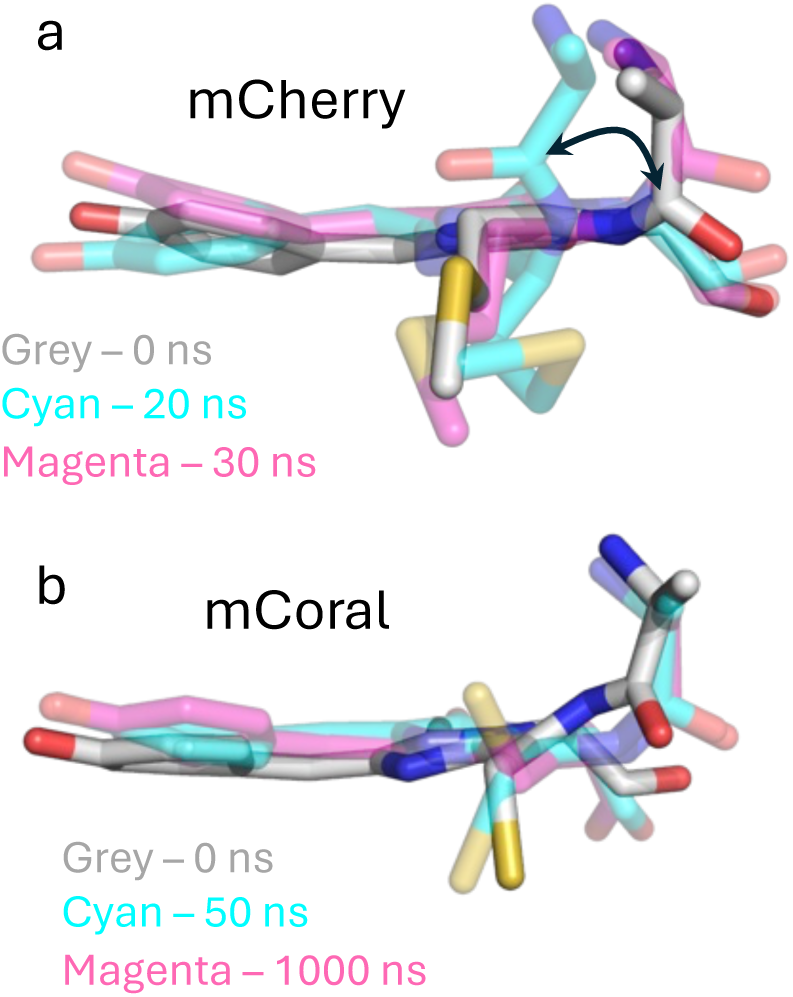
Different chromophore configurations of (a) mCherry and (b) mCoral extracted from 1000 ns simulations outlined on the figure. The structure for both were extract from the run 2 of the three 1000 ns MD simulations.

To mimic the experimentally observed solvated chromophore, we retain the original crystallographic waters during our simulations. To assess the impact of these water molecules, we rerun the MD simulations for mCherry starting with the protein only (no crystallographic waters) allowing de novo solvation, including in and around the chromophore. Chromophore switching was observed in one 500 ns simulation whereas switching was relatively rare in the other two 500 ns production runs (Figure S8). This suggests that the local water placement in mCherry could be influencing the exchange of the two F65 carbonyl oxygen states.

### 2.5 Chromophore associated water dynamics

We next looked at residency times and exit points of water molecules associated with the chromophore. The structurally conserved water molecule (W1 in Figure 4c) that H-bonds with the CRO phenol group typically has a low residency time in other FPs ^43^ and this also appear to be the case here (Table 3). The water does reside longer in mCoral (average 6.2 ns ± 1.9) than mCherry (average 0.6 ns ± 0.3) but still rapidly exchanges with the bulk solvent. For mCherry, the local exit point for the W1 water is between the bulge interrupted strand S7 and strand S10. Indeed, this is the common exit point region for all the CRO associated waters in mCherry (Reg 1 and 2 in Table 3; Figure 7). In contrast, the water exit points vary for mCoral (Figure 7b). The CRO phenol associated water can exit either side of the strand 7 bulge region encompassing either strand S10 (regs 1 or 2) or strand S8 (reg 3). However, the most common water exit point is reg 4 directly opposite the new thiol group which is between strands S3 and S11 (Figure 7b and Table 3) and includes the new W_M66C_ water molecule. Indeed, while we hypothesised the importance of W_M66C_ in maintaining the chromophore structure, its residency time is relatively short (1.7 ns ± 1.7; Table 3).

**Figure 7.**
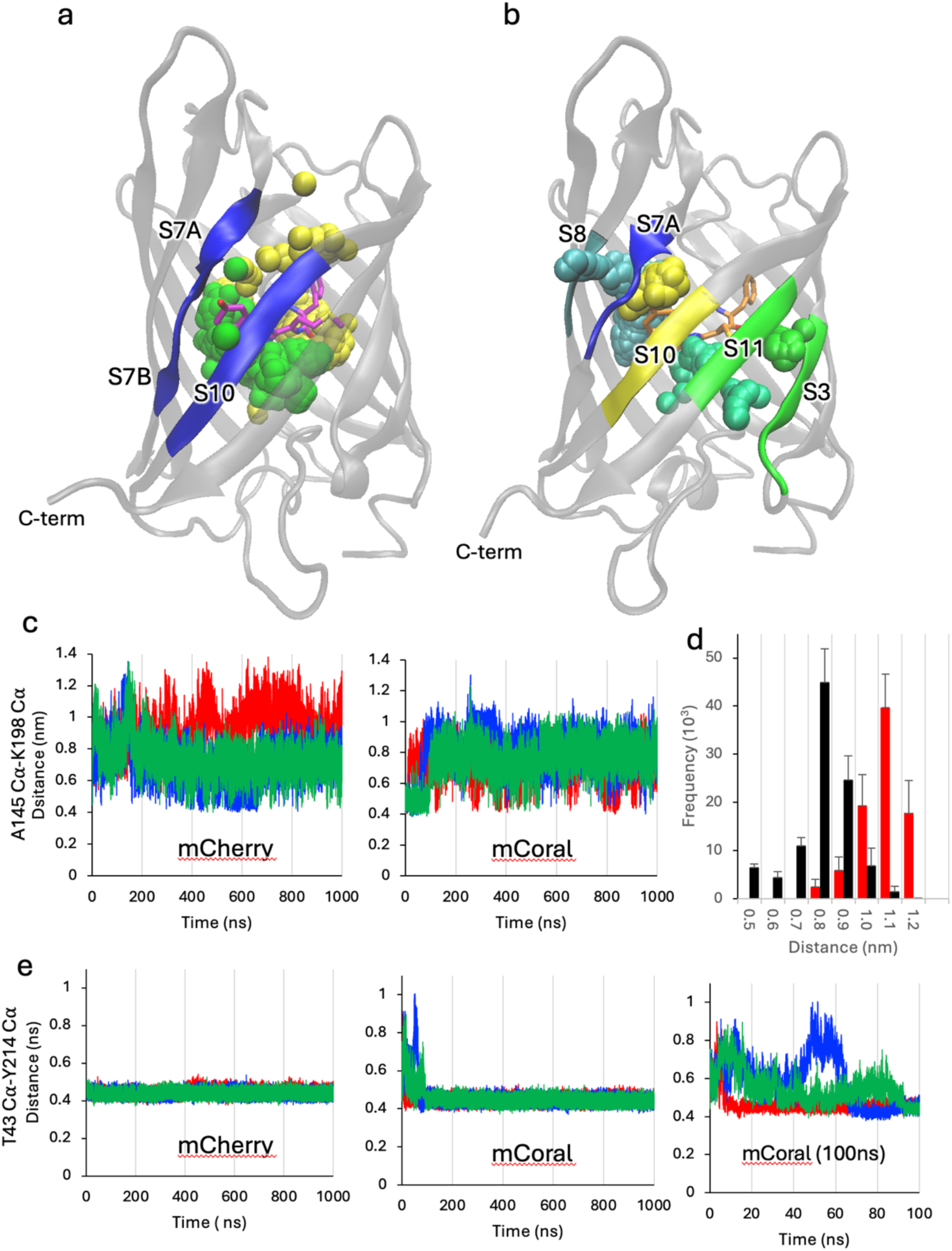
Chromophore associated water dynamics. Representative water exits points for (a) mCherry and (b) mCoral. The different sphere colours represent different water molecular paths. (c) S7-S10 Interstrand distances represented by the A145-K198 Cα distance. Green, blue and red represent the data from run 1, run 2 and run 3, respectively. (d) Distance distribution for mCherry (red) and mCoral (black) with bin sizes of 0.1 nm. (e) S3-S11 interstrand distances distances represented by the T43-Y214 Cα distance. Green, blue and red represent the data from run 1, run 2 and run 3, respectively. The right most panel is a zoom of the first 100 ns for the mCoral simulations.

**Table 3.**
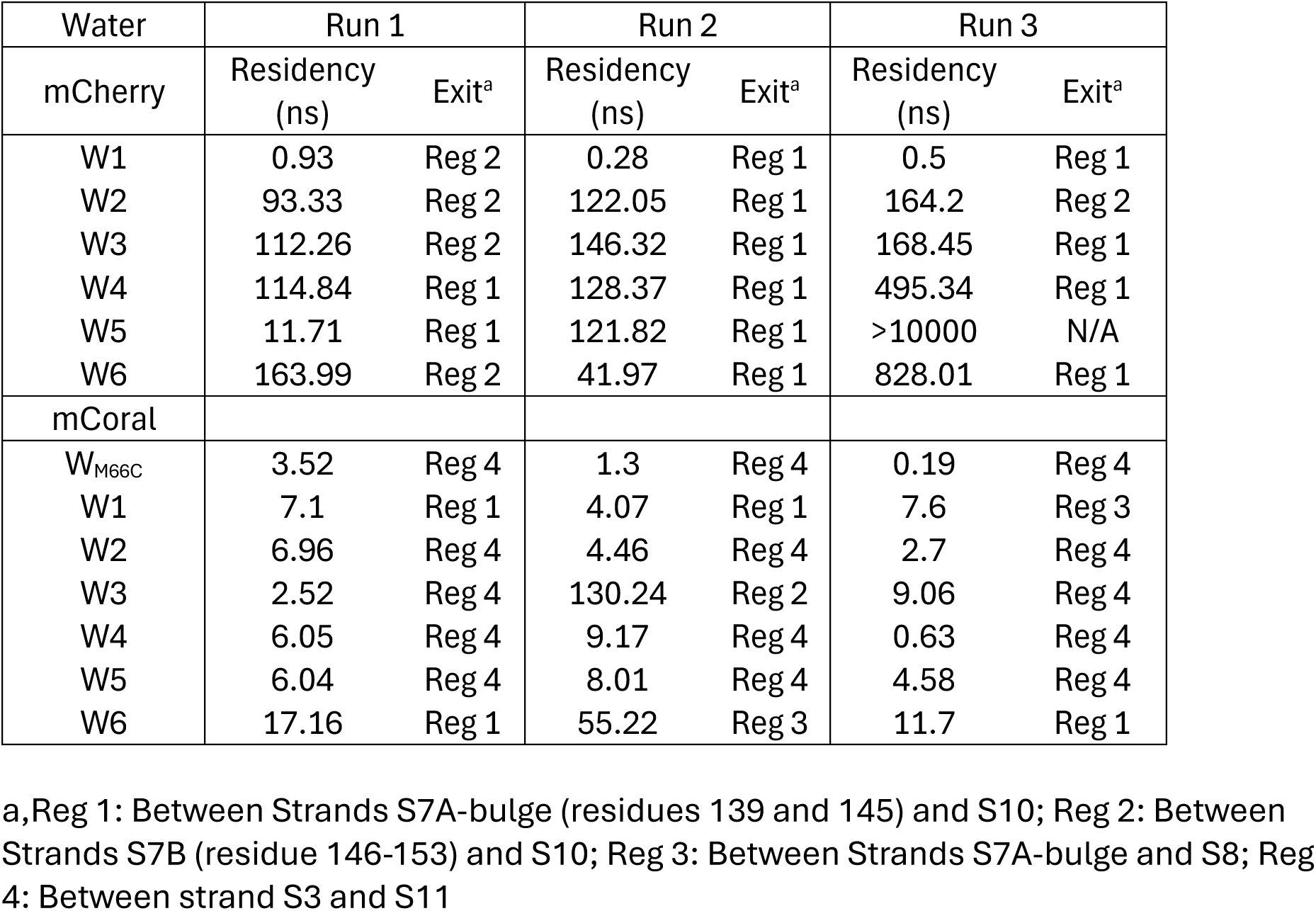
Water residency times and exit points.

| Water | Run 1 |  | Run 2 |  | Run 3 |  |
| --- | --- | --- | --- | --- | --- | --- |
| mCherry | Residency (ns) | Exit<sup>a</sup> | Residency (ns) | Exit<sup>a</sup> | Residency (ns) | Exit<sup>a</sup> |
| W1 | 0.93 | Reg 2 | 0.28 | Reg 1 | 0.5 | Reg 1 |
| W2 | 93.33 | Reg 2 | 122.05 | Reg 1 | 164.2 | Reg 2 |
| W3 | 112.26 | Reg 2 | 146.32 | Reg 1 | 168.45 | Reg 1 |
| W4 | 114.84 | Reg 1 | 128.37 | Reg 1 | 495.34 | Reg 1 |
| W5 | 11.71 | Reg 1 | 121.82 | Reg 1 | >10000 | N/A |
| W6 | 163.99 | Reg 2 | 41.97 | Reg 1 | 828.01 | Reg 1 |
| mCoral |  |  |  |  |  |  |
| W<sub>M66C</sub> | 3.52 | Reg 4 | 1.3 | Reg 4 | 0.19 | Reg 4 |
| W1 | 7.1 | Reg 1 | 4.07 | Reg 1 | 7.6 | Reg 3 |
| W2 | 6.96 | Reg 4 | 4.46 | Reg 4 | 2.7 | Reg 4 |
| W3 | 2.52 | Reg 4 | 130.24 | Reg 2 | 9.06 | Reg 4 |
| W4 | 6.05 | Reg 4 | 9.17 | Reg 4 | 0.63 | Reg 4 |
| W5 | 6.04 | Reg 4 | 8.01 | Reg 4 | 4.58 | Reg 4 |
| W6 | 17.16 | Reg 1 | 55.22 | Reg 3 | 11.7 | Reg 1 |

a, Reg 1: Between Strands S7A-bulge (residues 139 and 145) and S10; Reg 2: Between Strands S7B (residue 146-153) and S10; Reg 3: Between Strands S7A-bulge and S8; Reg 4: Between strand S3 and S11

While chromophore associated water residency does vary (with all but one exchanging with the bulk solvent), the lost waters are generally replaced by additional waters. This is highlighted by the persistency of H-bonds formed between the chromophore and water (Figure 5f). It still needs to be determined if water exchange rates are an important determinant FP photophysical properties but there is evidence that suggests it does ^43–46^.

To assess why the two proteins, have different dominant water exit points, we looked at local inter-strand distances. The exit point comprising strands S7 (regions S7A and S7B linked by a short bulge) and S10 does appear to be consistently shorter in mCoral compared to mCherry (Figure 7c and d). This region is a known water exit point in other FPs such as GFP ^43,44^. The most frequently observed distance is ∼0.8 nm for mCoral compared to ∼1.1 nm for mCherry (Figure 7d). It has been suggested previously that lower distance fluctuation between strands S7 and S10 plays a role in improving photobleaching resistance in the mCherry-XL variant (compared to mCherry) ^46^. Given that mCoral also has shorter distance fluctuations in this region (Figure 4c-d), this may also explain its improved resistance to H_2_O_2_.

Strand S10 is linked to S11, which along with S3 forms a major water exit point for mCoral. In both proteins, the distance between strands S3 and S11 remains relatively stable (Figure 7e). However, the T43 (S3) and Y214 (S11) Cα distance in mCoral fluctuates within the first 100 ns before it stabilises. It is during this period of flux that exchange of waters occur (Table 3), with no water exiting beyond 10 ns. Thus, local increased solvation on the introduction of the M66C mutation into mCherry does have repercussions in terms of local secondary structure stability. The new local solvent exit/entry point opposite the new thiol group may also contribute to mCorals increased susceptibility to changes in pH.

## 3. Conclusion

Fluorescent proteins have proved to be pivotal in our understanding of biological processes, with mCherry being one of the most utilised FPs in the red region of the visible spectrum. The simple mutation of methionine to cysteine in the chromophore to generate mCoral adds new features to mCherry. Given the role of pH in biology, ^47,48^ the pH responsive nature of mCoral across a broad range opens up the possibility of it being used to monitor changes in different cellular compartments. mCoral could also be used in place of mCherry in environments where ROS levels are relatively high ^49^. While the M66C mutation has relatively little impact on protein structure, it does change two vital aspects: the water molecules local to the chromophore and the dynamics. Our long-time scale MD of mCherry reveals that the chromophore itself could be undergoing conformational flux, switching between two distinct configurations. Such switching does not appear to be happening in mCoral potentially due to a new water molecule that forms polar interactions with the carbonyl oxygen that undergoes the flip in mCherry. This has consequences in terms of future engineering of mCherry to improve its brightness through increasing quantum yield. The quantum yield of a FP is known to be influenced by its chromophore dynamics so engineering next generation brighter FPs in the red region should take chromophore dynamics into account. The same is true for water dynamics. Internal waters molecules close to the chromophore help shape FPs properties, with increased organisation likely playing a key role in improved brightness ^43,44,50,51^. The next step to understand how water dynamics are related to spectral properties. This in turn can help us establish rules to help predict how best to organise such waters through changes in the protein.

## 4. Methods and materials

### 4.1 Site directed mutagenesis

The mCherry encoding gene resident in the pBAD bacterial expression plasmid has been described previously ^50,52^. The M66C mutation was introduced by whole plasmid, inverse PCR using Q5 High Fidelity DNA polymerase kit (New England Biolabs) in combination with a forward (5’-GCTCCAAGGCCTACGTGAAG-3’) and mutagenic reverse (5’-CGTA**<u>GCA</u>**GAACTGAGGGGACA-3’; mutation site shown as bold and underlined) primers (synthesised by Integrated DNA Technologies) using the manufacture’s protocol. Plasmids were sequenced (EuroFins Genomics) to confirm the presence of the mutation.

### 4.2. Recombinant protein production

The pBAD plasmids encoding mCherry or mCoral (mCherry M66C) were chemically transformed into *E. coli* TOP™ cells (Invitrogen) and plated on LB agar plates supplemented with 50 μg/mL ampicillin. A single colony was used to inoculate a 5 mL overnight culture, which was then used to inoculate of 2xTY media supplemented with 50 μg/mL ampicillin. Cultures were left to grow at 37°C until they reached an OD_600_ of 0.6 when 0.2% (v/v) arabinose was added to induce expression. The culture was left to incubate overnight at 37° in a shaking incubator. Cells were harvested via centrifugation at 5000 x *g* for 20 minutes at 4°C. The cell pellet was resuspended in 50 mM Tris-HCl buffer (pH 8) and cells were lysed using a French press. The resulting lysate was clarified by centrifugation at 25000 x *g* for an hour at 4°C and the resulting supernatant was collected. Clarified cell lysate was loaded onto a 5 ml His-trap HP nickel affinity column (Cytiva) equilibrated in wash buffer (50 mM Tris-HCl, 10 mM imidazole pH 8.0) for Nickel-affinity chromatography. Bound protein was eluted by washing the column with 500 mM imidazole (pH 8.0). Pooled protein samples were then subjected to size exclusion chromatography (SEC) using a HiLoad 26/600 Superdex 200 column that was equilibrated with 50 mM Tris- HCl buffer (pH8.0). Purity of the proteins was then checked by SDS-PAGE.

### 4.3. Spectral analysis

Absorbance and fluorescence measurements were performed using Cary 60 UV-Vis spectrophotometer (Agilent technologies) and Varian Carry Eclipse Fluorescence spectrophotometer (Agilent Technologies), respectively. Absorbance spectra were recorded using a 1 cm path-length quartz cuvette. Protein fluorescence spectra were recorded using a 5×5 mm quartz cuvette and data was collected with a 5 nm slit width at a rate of 600 nm/min. Each protein was excited at their respective excitation maximum. Spectra were recorded at concentrations of 2.5 μM or 5 μΜ. Molar absorbance for mCherry at its λ_max_ was calculated to be similar to that reported in FPBase (https://www.fpbase.org/protein/mcherry/) and used to generate a molar absorbance (ε) value at 280 nm ε_280_. The ε_280_ for mCherry was then used to calibrate spectra for mCoral (as there are no additional aromatics present) to determine ε. Quantum yield of mCoral was determined as described previously ^50^ using mCherry as the standard. The pH profile analysis was performed as described previously ^43^ using 5 μM of protein at pH 4.5, 5.0, 5.5 (all 100 mM acetate buffer), 6.0 (100 mM KH_2_PO_4_), pH 7.0 (100 mM HEPES), 8.0 (50 mM Tris-HCl), 9.0, 10.0 (both 100 mM glycine-NaOH) 11.0 (100 mM Na_2_HPO_4_- NaOH). Tolerance to H_2_O_2_ was determined using 2.5 μM of protein mixed with increasing concentrations of H_2_O_2_ ranging from 0.01% (v/v) to 5% (v/v). Absorbance and fluorescence were recorded at various time intervals.

### 4.4 Mass spectrometry analysis

Pure mCherry and mCoral were diluted to 10 μM and analysed by the Mass Spectrometry facility in the School of Chemistry, Cardiff University. The sample was subject to liquid chromatography-mass spectrometry (LCMS) using a Waters Acquity UPLC/Synapt G2-Si QTOF mass spectrometer. For the H_2_O_2_ treated sample, H_2_O_2_ was added just prior to the LC step.

### 4.5. DFT analysis

All DFT analysis employed the Orca 6.1.0 package ^53^. Geometry optimisation used the PBE-D3BJ^54,55^/def2-SVP ^56^ level without any constraint, and confirmed as minima using harmonic frequency calculation. Absorption spectra and frontier molecular orbitals were then calculated at optimal geometry at PBE0^57^/def2-TZVP level in CPCM ^58^ model of aqueous solution. Orbital plots were obtained using Avogadro v 1.2.0.

### 4.6. mCoral structure determination

Crystal trials of purified protein samples (∼10mg/ml) were set up at Diamond Light Source (DLS), Harwell, UK. Crystal formation was screened using sitting drop vapour diffusion across a wide variety of conditions as described by the BSC, PACT, premierTM, JCSG-plusTM, SG1TM and MorpheusR crystallisation screens (Molecular Dimensions). Drops were set up with various protein:crystallisation buffer ratios including 1:1, 1:2 and 2:1 using a mosquito (SPT) and left in Formulatrix imagers at VMXi to incubate at room temperature until crystals formed. Data was collected at VMXi and at I03. The process of crystallisation and data acquisition was automated at DLS. Crystals were formed under 0.2 M magnesium chloride, 0.1 M Tris buffer, pH 8, glycerol 10 % v/v, 25% PEG 6-10kD and passed through the I03 DLS beamline. Following data acquisition, diffraction data were integrated and reduced using the XIA2 package ^59^. The structure was solved by molecular replacement using PHASER using the structure of mCherry (PDB 2H5Q was the search object using a process described previously ^60,61^. PHASER confirmed there was only one copy in the asymmetric unit producing a large Log likelihood score of 6303 and TFZ (Twin Fraction Zero refinement) of 26, indicating high confidence in the result. The model was adjusted to match the correct sequences, and to fit it into the electron density map using COOT ^62^ and then refined with TLS (Translation/Libration/Screw-rotation) parameters using RefMac5 ^63,64^. All programs were accessed via the CCP4 package ^65^ (https://www.ccp4.ac.uk).

### 4.7. Molecular dynamics simulations

Molecular Dynamics (MD) simulations were performed on the Supercomputing Wales HAWK server (project code scw1631). Molecular dynamics were performed using GROMACS ^66,67^ CHARMM27 forcefield ^68,69^ modified to contain constraints for the mCherry and mCoral chromophore in the CRO-O^-^ form. The forcefield parameters are available on request. The protein was then placed centrally in a cubic box at least 1 nm from the box edge applying periodic boundary conditions. The protein system was then solvated with water molecules (TIP3P) and total charge balanced to zero with Na^+^ ions.

The protein was then energy minimized to below 1000 kJ mol^−1^ nm^−1^ with an energy step size of 0.01 over a maximum of 50000 steps. The system was then temperature and pressure equilibrated using the NVT (constant number of particles, volume and temperature) followed by NPT (number of particles, pressure and temperature) ensembles. MD production runs were then performed at 300 K, 1 atmosphere pressure for 1000 ns with a 2 fs time step integration. Each protein was subjected 3 independent production runs. The original crystallographic waters present in the crystal structure were included in the 1000 ns production runs. The MD process was repeated for mCherry with all crystallographic waters removed and added back de novo during the solvation step, with MD production runs of 3 x 500 ns. The protein in the trajectories was centred in the simulation box and dumps of individual trajectories were performed via the *trjconv* command. RMSD and RMSF calculations were performed using the *rms* and *rmsf* commands. Pairwise distances and hydrogen bonds were determined using the *pairdist* and *hbond* commands. The recommended hbond default parameters were used (https://manual.gromacs.org/current/onlinehelp/gmx-hbond.html).

## Author Contributions

A.Z. contributed to the conception of the project, generated the mCoral mutant, produced protein, undertook the spectral analysis, contributed to structure determination and contributed towards the molecular dynamics. O.Z. contributed towards the molecular dynamics analysis and some aspects of mCherry characterisation. D.V. contributed to structure determination. P.J.R. contributed to structure determination. H.M. contributed to protein crystallisation and X-ray diffraction measurements. J.A.P. undertook DFT analysis. G.E.M contributed towards molecular dynamics data generation and analysis and help direct the project. D.D.J contributed to project conception and directed the project, contributed to general data analysis and molecular dynamics analysis. All authors contributed to the writing of the paper and analysing data.

## Institutional Review Board Statement

Not applicable.

## Informed Consent Statement

Not applicable.

## Data Availability Statement

The mCoral structure and associated data has been deposited in the Protein Data Bank under PDB ID 9TAD. All spectroscopic data will be made available via FigShare (To be released on publication).

## Acknowledgments

The authors would like to thank the Cardiff School of Biosciences Protein Technology Hub for helping with the production and analysis of proteins. We would also like to thank Advanced Research Computing at Cardiff (ARCCA) facility for supporting our molecular dynamics work. We would also like to thank Diamond Light Source, especially the VXMi facility and its staff for the invaluable service they provide. We would like to thank the Crystallization Facility at Harwell for access and support.

