## Supplementary material for "Structure, function and dynamics of mCoral, a pH responsive engineered variant of the mCherry fluorescent protein with improved hydrogen peroxide tolerance": SI Figures and Tables.

**Supplementary Information.**

**Supplementary Table S1**. Data collection and refinement statistics for mCoral

| **PDB Entry** | |
|---|---|
| **Data Collection*** | |
| Diamond Beamline | I03 |
| Date | 2021-09-30 |
| Wavelength | 0.81532 |
| **Crystal Data (figures in brackets refer to outer resolution shell)** | |
| *a,b,c* (Å) | 48.760, 43.182, 63.095 |
| $\alpha,\beta,\gamma$ | 90.0, 90.114.91, 90.0 |
| Space group | P 1 $2_1$ 2 |
| Resolution (Å) | 2.04 – 45.46 |
| Outer shell | 2.04 – 2.10 |
| *R*-merge (%) | 9.1 (46.7) |
| *R-pim (%)* | 9.1 (46.7) |
| *R*-meas (%) | 12.8 (66.1) |
| CC1/2 | 0.980 (0.413) |
| I / σ(I) | 20.5 (4.7) |
| Completeness (%) | 97.3 (98.3) |
| Multiplicity | 1.8 (1.8) |
| Total Measurements | 27,152 (2,107) |
| Unique Reflections | 14,902 (1,149) |
| Wilson B-factor(Å$^2$) | 20.2 |
| **Refinement Statistics** | |
| Refined atoms | 1,853 |
| Protein atoms | 1,761 |
| Non-protein atoms | 5 |
| Water molecules | 87 |
| R-work reflections | 13,309 |
| R-free reflections | 1,376 |
| R-work/R-free (%) | 16.98 / 21.06 |
| **rms deviations (ML target in brackets)** | |
| Bond lengths (Å) | 0.012 (0.013) |
| Bond Angles (°) | 1.553 (1.661) |
| [1]Coordinate error | 0.111 |
| Mean B value (Å$^2$) | 23.4 |
| **Ramachandran Statistics (PDB Validation)** | |
| Favoured/allowed/Outliers | 208 / 9 / 0 |
| % | 95.9 / 4.1 / 0.0 |

* One crystal was used for determining each structure.
[1] Coordinate Estimated Standard Uncertainty in (Å), calculated based on maximum likelihood statistics.

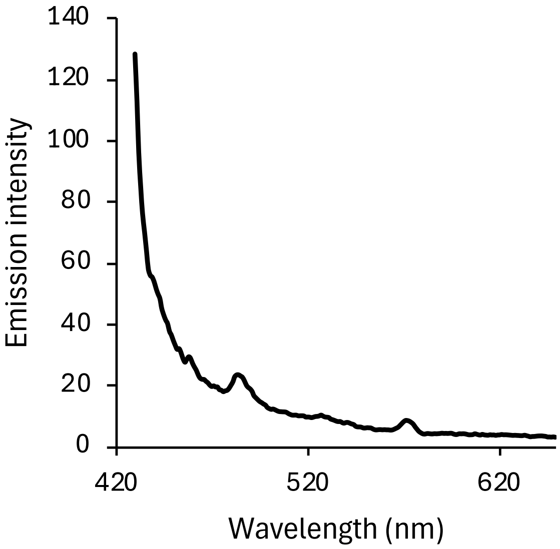


**Supplementary Figure S1**. The emission spectra of mCoral at pH 4.5 on excitation at the $\lambda_{max}$ for that pH (420 nm).

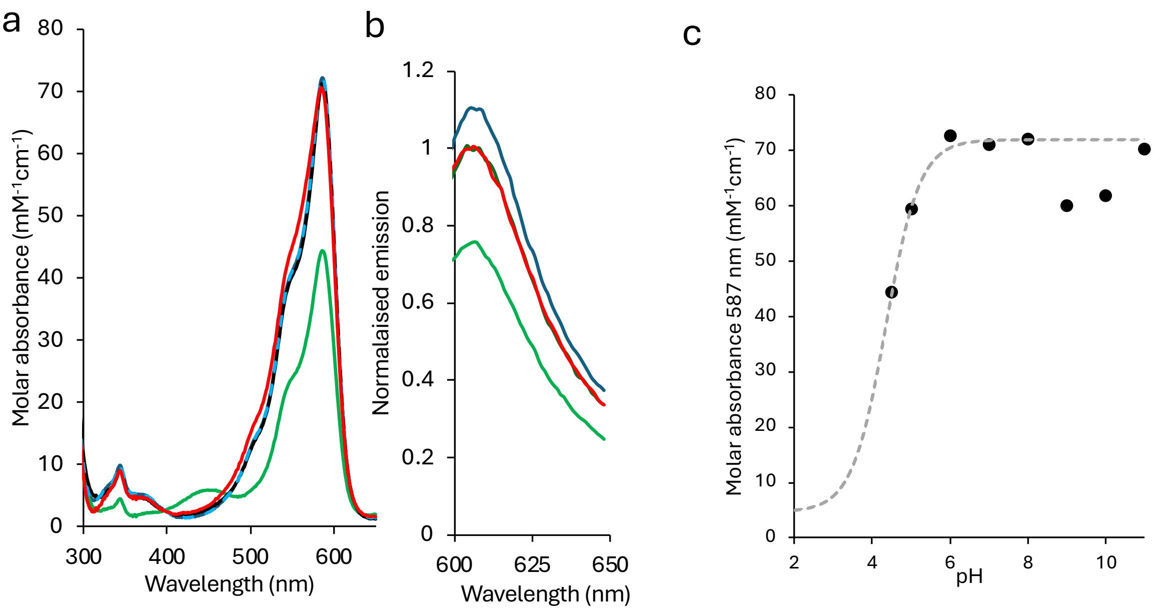


**Supplementary Figure S2**. The effect of pH on the spectral properties of mCherry. Change in (a) absorbance spectra and (b) emission spectra (on excitation at $\lambda_{max}$) at pH 4.5 (green), pH 7 (black), pH 8 (dashed blue line) and pH 11 (red). (c) Plot of change in 587 nm molar absorbance against pH fitted to a single transition sigmoidal curve according to the Henderson-Hasselbach equation.

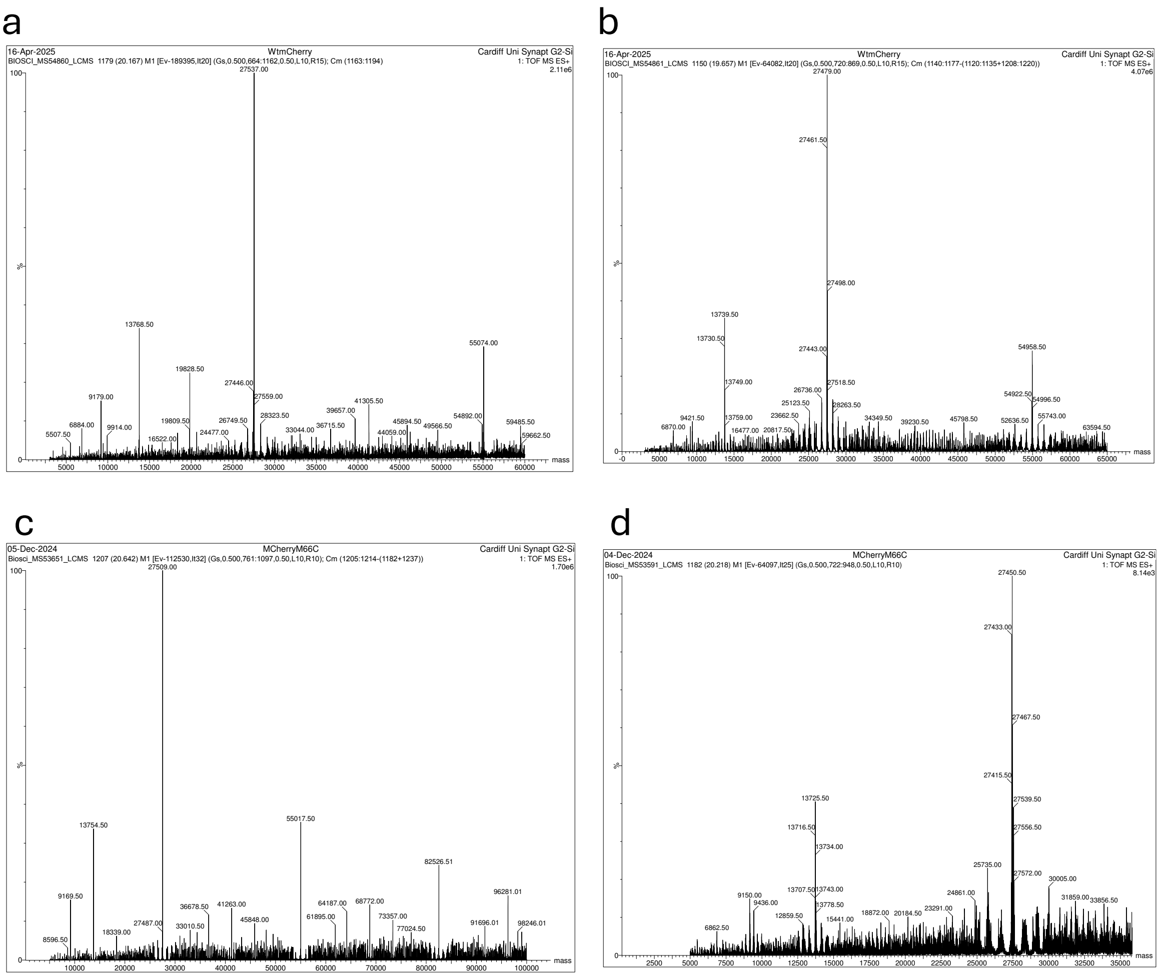


**Supplementary Figure S3.** Mass spectra of mCherry before (a) and after (d) addition of 0.1 % (v/v) $H_2O_2$, and of mCoral before (c) and after (d) addition of 0.1 % (v/v) $H_2O_2$.

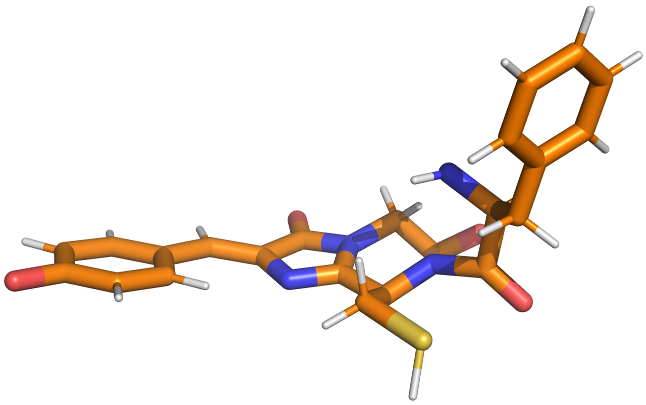

**Supplementary Figure S4.** The mCoral chromophore unit used in MD simulations.

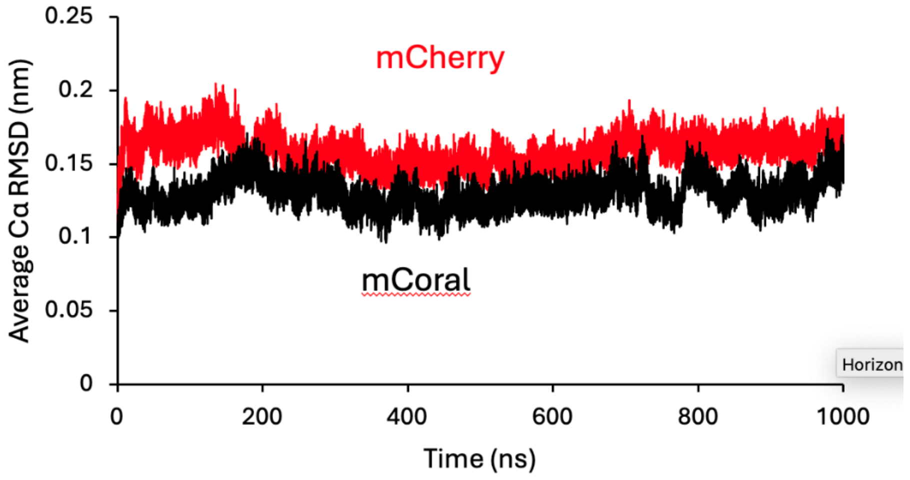


**Supplementary Figure S5.** Average Cα RMSD over 3 x 1000 ns MD runs for mCherry (red) and mCoral (black).

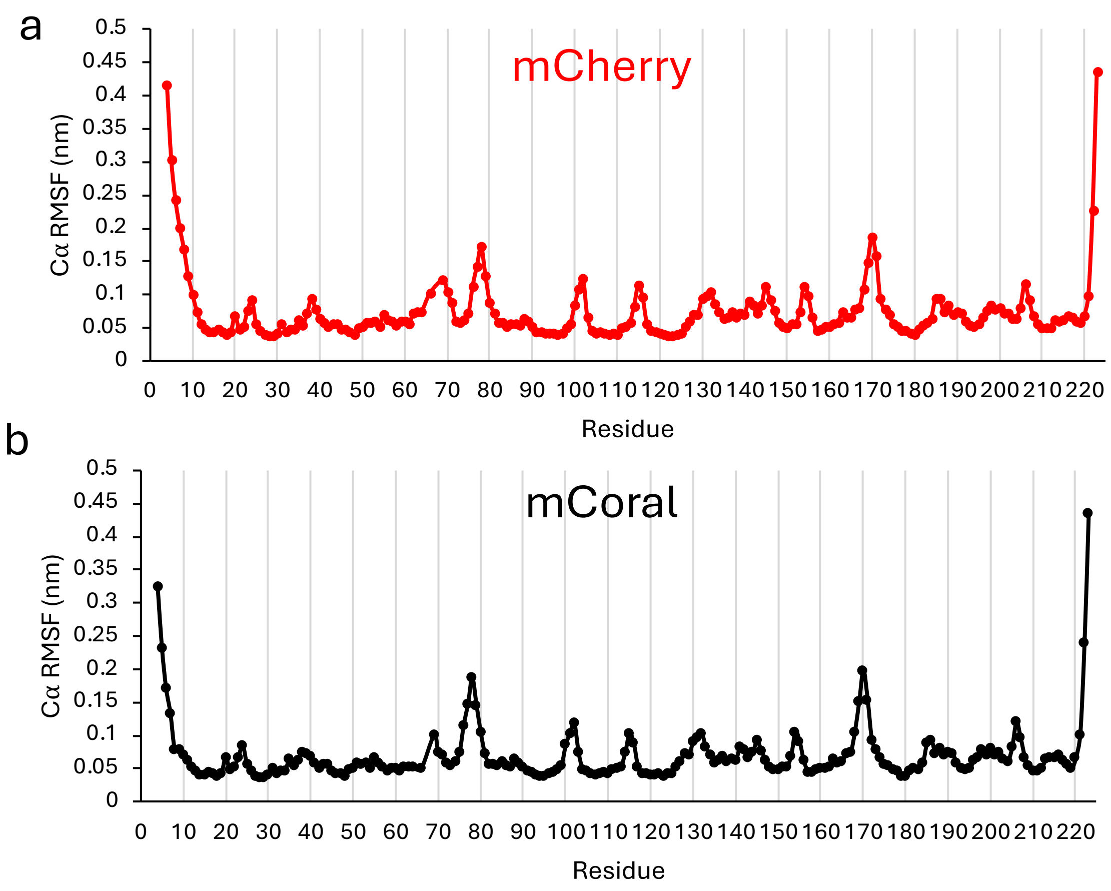


**Supplementary Figure S6.** Average Cα RMSF over 3 x 1000 ns MD runs for (a) mCherry and (b) mCoral.

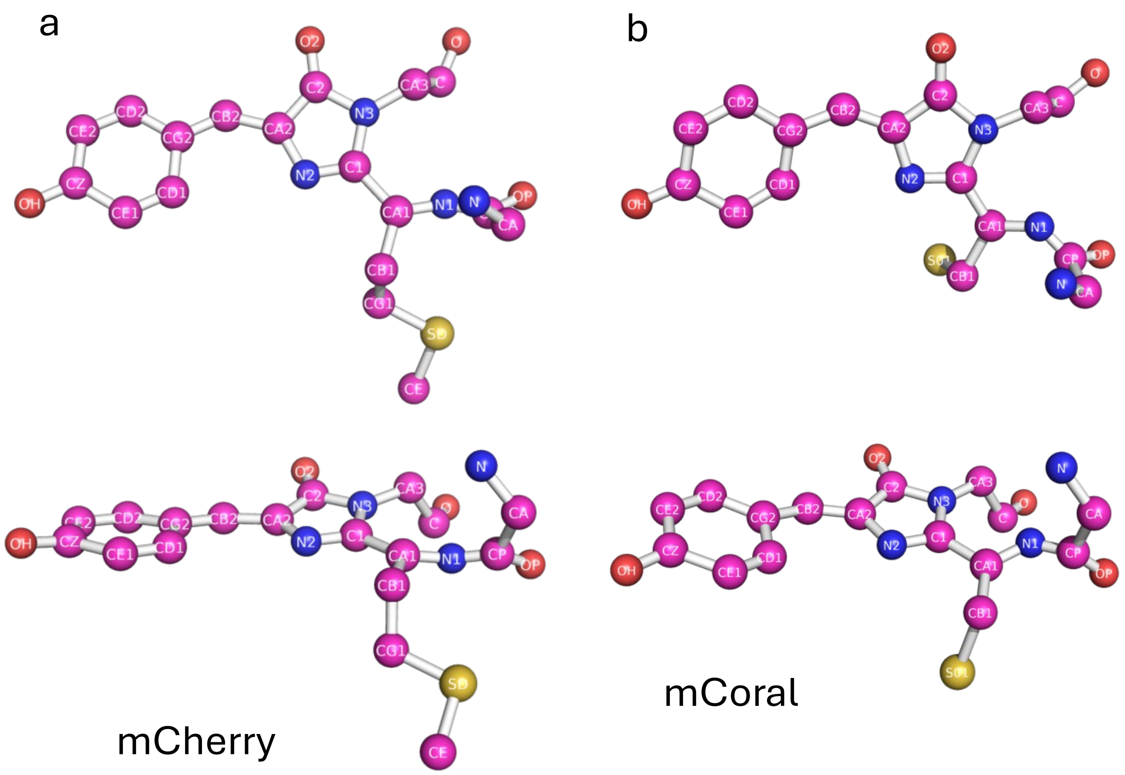


**Supplementary Figure S7.** CRO atom nomenclature for (a) mCherry and (b) mCoral.

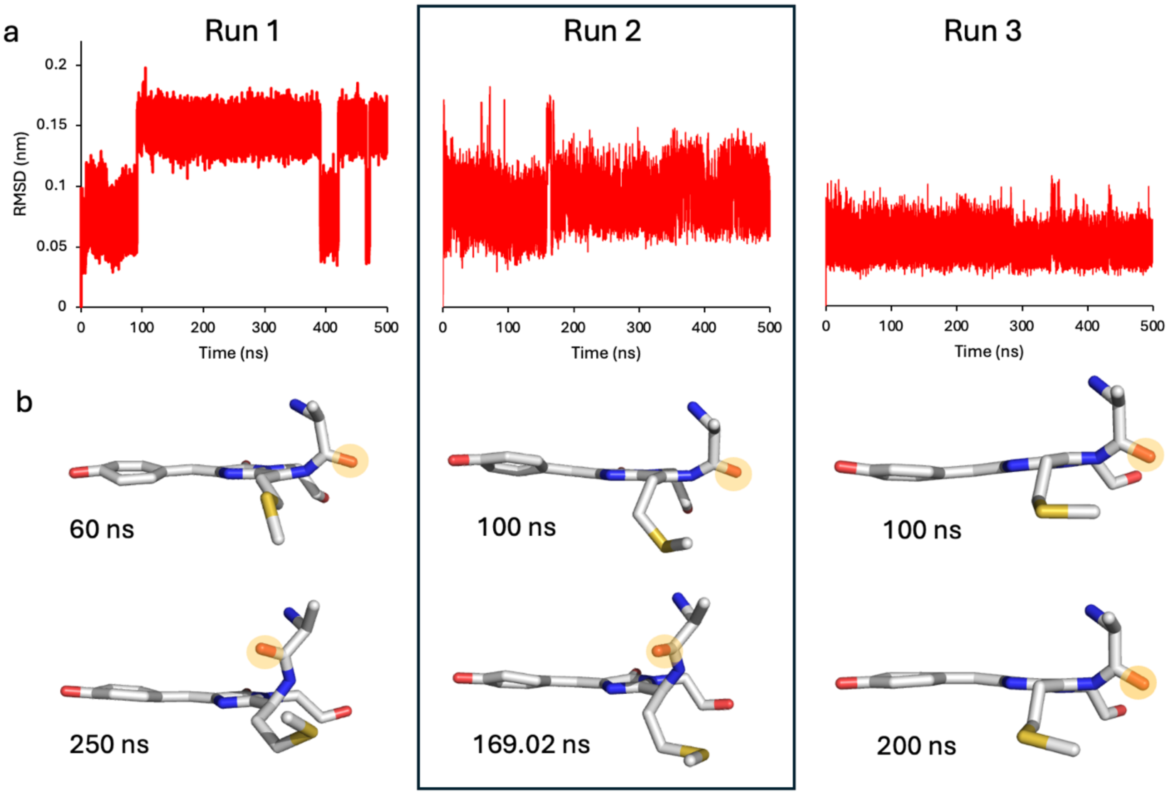


**Supplementary Figure S8.** MD simulations of mCherry with de novo added water. (a) The CRO (all atoms) RMSD profile over the 3 runs. (b) Different chromophore configurations of mCherry extracted from the 500 ns simulations at time points outlined on the figure. The orange circle highlights the F65 carbonyl oxygen.
